# Elevation-related climate trends dominate fungal co-occurrence patterns on Mt. Norikura, Japan

**DOI:** 10.1101/2021.01.25.428196

**Authors:** Ying Yang, Yu Shi, Dorsaf Kerfahi, Matthew C Ogwu, Jianjun Wang, Ke Dong, Koichi Takahashi, Itumeleng Moroenyane, Jonathan M. Adams

## Abstract

Although many studies have explored patterns of fungal community diversity and composition along various environmental gradients, the trends of co-occurrence networks across similar gradients remain elusive. Here, we constructed co-occurrence networks for fungal community along a 2300 m elevation gradient on Mt Norikura, Japan, hypothesizing a progressive decline in network connectivity with elevation due to reduced niche differentiation caused by declining temperature and ecosystem productivity. Results agreed broadly with predictions, with an overall decline in network connectivity with elevation for all fungi and the high abundance phyla. However, trends were not uniform with elevation, most decline in connectivity occurred between 700 m and 1500 m elevation, remaining relatively stable above this. Temperature and precipitation dominated variation in network properties, with lower mean annual temperature (MAT) and higher mean annual precipitation (MAP) at higher elevations giving less network connectivity, largely through indirect effects on soil properties. Among keystone taxa that played crucial roles in network structure, the variation in abundance along the elevation gradient was also controlled by climate and also pH. Our findings point to a major role of climate gradients in mid-latitude mountain areas in controlling network connectivity. Given the importance of the orographic precipitation effect, microbial community trends seen along elevation gradients might not be mirrored by those seen along latitudinal temperature gradients.

**Importance:** Although many studies have explored patterns of fungal community diversity and composition along various environmental gradients, it is unclear how the topological structure of co-occurrence networks shifts across environmental gradients. In this study, we found that the connectivity of the fungal community decreased with increasing elevation, and that climate was the dominant factor regulating co-occurrence patterns, apparently acting indirectly through soil characteristics. Assemblages of keystone taxa playing crucial roles in network structure varied along the elevation gradient and were also largely controlled by climate. Our results provide insight into the shift of soil fungal community co-occurrence structure along elevational gradients, and possible driving mechanisms behind this.

**Graphic abstract:** 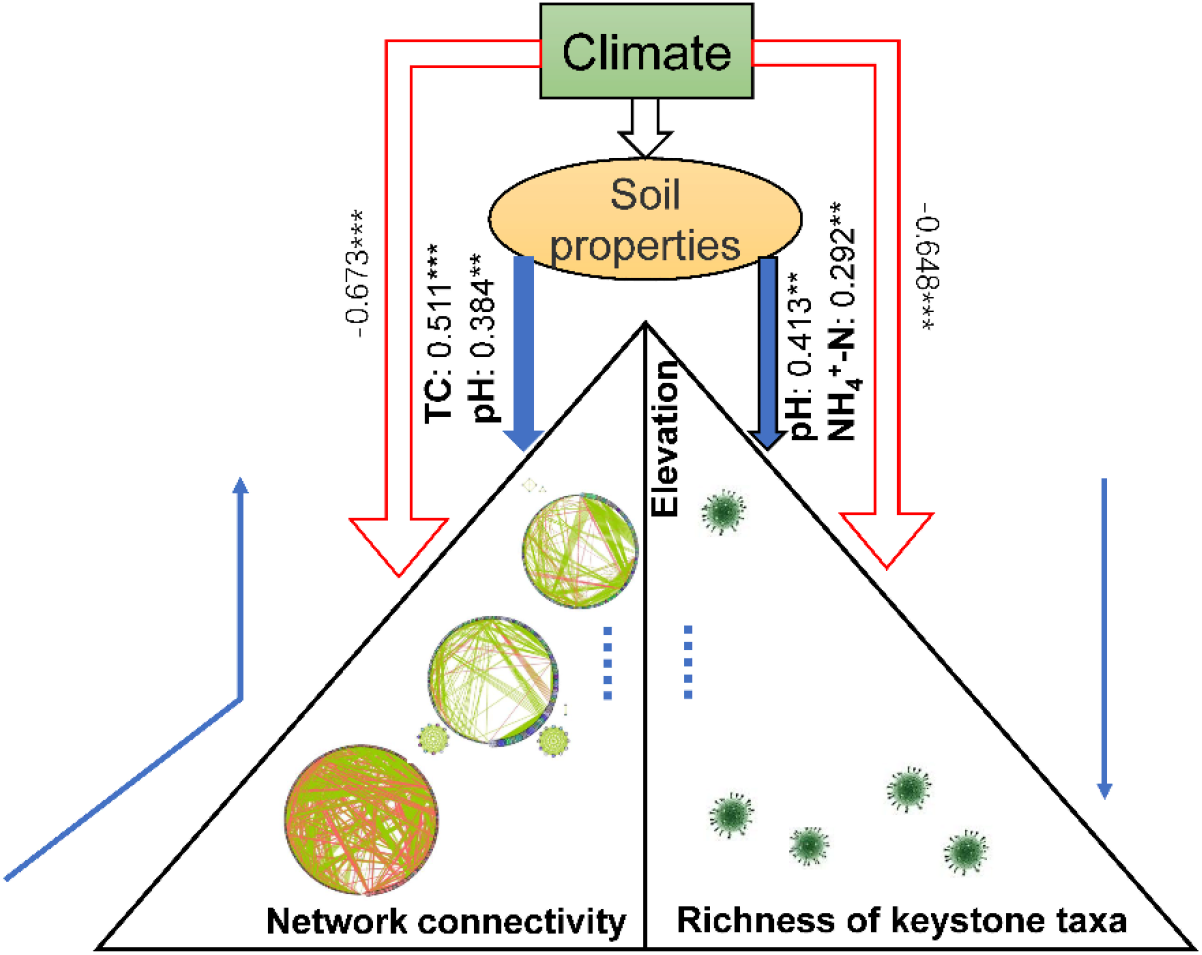

## 1. Introduction

In the soil microbial community, fungi play a critical role because they promote nutrient cycling in soil ecosystem mainly as decomposers of plant litters (1). Additionally, many fungal taxa can form a mutualistic relationship with plants to affect their nutrient and water absorption, and protect plants from the influence of biotic and abiotic stress (2, 3). Thus, understanding the structuring of soil fungal communities, and by the possible implications for ecosystem stability, cycle and maintenance, is one of the most important goals in ecology.

Interspecies interactions within fungal communities are poorly understood but potentially important (4). Many microorganisms interact with one another directly through interspecies interactions (e.g., through mutualistic and competitive interactions), and also interact in common with larger host organisms, bringing about indirect associations with other microorganism species (5). Such interactions often cannot be observed directly, but can be inferred through microbial co-occurrence network analyses utilizing high throughput metagenetic data (6; 7). Network analysis summarizes the number, frequency and identity of links between different species in the community (8). The types of interspecific association are usually classified into positive and negative, based on increased or decreased likelihood of cooccurrence, and should not be confused with positive and negative effects on fitness. For example, positively associated co-occurrence can result from co-colonization, niche overlap and/or cross-feeding, while negative cooccurrence associations between taxa may result from the present day or past evolutionary effects of competitive exclusion, where species have discrete specialised niches and amensalism (9, 10).

Topologically, different OTUs (nodes) play distinct roles in the network (11). keystone taxa are defined as those that occupy a considerable role in community structure and integrity, and their influence is independent of abundance (12). These taxa play a unique and crucial role in the microbial community, as their removal can result in dramatic shift in the structure and functioning of a microbiome (13, 14, 15). For example, *Pseudomonadaceae* have been found to contribute to the natural suppressiveness of soils against the fungal pathogens in the rhizosphere (16). *Chaetomium*, *Cephalotheca* and *Fusarium* in fungi had strong positive association with organic matter decomposition rate, indicating their importance in C turnover (17). However, few studies have investigated the keystone taxa of fungal communities in mid-altitude mountain areas, let alone the environmental factors that regulate the abundance and distribution of these.

In fungal community studies, various trends in network structure have been noted along ecological gradients or between different types of environment (18). Xiao et al. (19) constructed networks for soil fungal communities of restored and natural salt marshes, finding that restored (desalinized) marsh had a more stable network compared to that of natural salt marsh with halophytic plant species. Guo et al. (20) investigated the co-occurrence patterns of 140 fungal communities along a soil fertility gradient, finding that fungal networks were larger and showed higher connectivity as well as greater potential for inter-module connection in more fertile soils. On a much broader geographical scale, Hu et al. (21) carried out an investigation on forest soil across five climate zones in China, revealing there was a hump-shaped pattern of interaction strength between fungal species from high-latitude towards low-latitude. So far, across the literature, there is a complex picture on the factors affecting co-occurrence network structure. Ma et al. (22) compared the network structure of fungal and bacterial communities in soil systems in different regions along a latitudinal gradient in eastern China, finding that more northerly parts of China had greater network complexity than those further south. They also suggested that this trend may occur due to greater precipitation in the south of China bringing about more chemically uniform weathered soils, with fewer opportunities for microhabitat niche differentiation and less evolutionary selection for evolving precise interactions in different community types. However, there is a pressing need to study how network patterns in soil biota vary among other temperature gradients, to understand whether the same relationship between network complexity and climate holds true elsewhere, and whether the sort of ecological mechanisms Ma et al proposed – or other additional mechanisms – might hold true. Testing different systems with different combinations of environmental conditions can help to disentangle how community structure varies and whatever underlying mechanisms are at work. Abiotic factors such as soil pH, photosynthetic carbon availability, precipitation, spatial distance between sites may be the key factors shaping fungal co-occurrence networks (22, 23, 24, 25), and microbial phylogeny has also been found to affect network patterns (26).

Elevational gradients are characterized by drastic shifts in abiotic and biotic factors – largely driven by climate - over short geographic distances (27, 28). As such they may offer opportunities for discerning the drivers of broader scale biogeographic patterns in community structure and processes. Nevertheless, elevation gradients have so far been relatively little studied in terms of microbial community network patterns. Qian et al. (28) focused on the elevational effects on the phyllosphere fungal assemblages in a single tree species, demonstrating that phyllosphere fungal networks showed reduced connectivity with increasing elevation. To our knowledge, soil fungal network structure has not been studied from the perspective of elevation.

Here, in response to the lack of studies along elevation gradients, we chose as our study focus Mt Norikura in central Japan. Norikura is an extinct volcano, last active about 18,000 years ago, which provides a broad climate gradient between its lower elevations around 700m and its summit at around 3,000m (29). We hypothesized that (1) the network connectivity of total fungal community and of the most abundant phyla would decrease with increasing elevation, with mean annual temperature dominating the process. This would principally be as a result of reduced energy flow through the ecosystem due to decreasing primary productivity at lower temperatures, preventing niche specialisation due to resources being less abundant and less stable. (2) We also hypothesized that the diversity of keystone taxa would shift dramatically along the elevational gradients, with fewer keystone taxa playing a role in the network, due to the prevalence of more generalized interactions.

## 2. Results

### 2.1 Elevation patterns for environmental parameters

Climate-model estimated climate and measured soil environmental variables varied considerably between different elevations (Figure S1). Several variables exhibited distinct elevational gradients. For example, MAT and NO_3_^−^-N decreased with increasing elevation, whereas pH showed a U-shaped trend, and remaining environmental variables-elevation are unimodal.

### 2.2 Meta-community co-occurrence network and sub-networks

To understand the role of biotic interaction in community assembly, the co-occurrence network analysis of fungi (at the OTU level) on Mt Norikura was constructed based on significant correlations. For the whole fungal community, the meta-community co-occurrence network captured 6725 associations (edges) among 358 OTUs (nodes), with 98.43% positive edges and 1.57% negative edges (Figure S2). The degrees for fungi were distributed according to power-law distributions (Figure S3 at [https://doi.org/10.6084/m9.figshare.13625609.v1]), which indicated a scale-free network structure and a non-random co-occurrence pattern, meaning that most OTUs had low-degree values, and only a few hub nodes had high-degree values (Ma, B. et al., 2016). The majority of fungal sequences belonged to the phyla Ascomycota (relative abundance 65.1%), Basidiomycota (20.4%) and Zygomycota (0.09%), the associations were mainly observed among order *Helotiales*, *Agaricales, Eurotiales, Sordariales, Chaetothyriales, Pleosporales* and *Hypocreales* (Table S1). Some topological properties commonly used in network analysis were calculated to describe the complex pattern of interrelationships between OTUs (30). The average network distance between all pairs of nodes (average path length) was 3.075 edges with a diameter (longest distance) of 11 edges. The clustering coefficient (that is, how nodes are embedded in their neighbourhood and, thus, the degree to which they tend to cluster together) was 0.777 and the modularity index was 0.235 (values >0.4 suggest that the network has a modular structure). The sub-network diagram of each elevational level is shown in Fig 1, network edge density increases with the elevation to 1500m, and above this does not change significantly with the elevation. Overall, the soil fungal community network was comprised of highly connected OTUs (in terms of edges per node) structured among densely connected groups of nodes (that is, modules) and forming a clustered topology, as expected for real-world networks that are more significantly clustered than random graphs. These structural properties offer the potential for ready comparisons among complex datasets from different ecosystem types, in order to explore how the general traits of a certain habitat type may influence the assembly of microbial communities.

**Fig 1.**
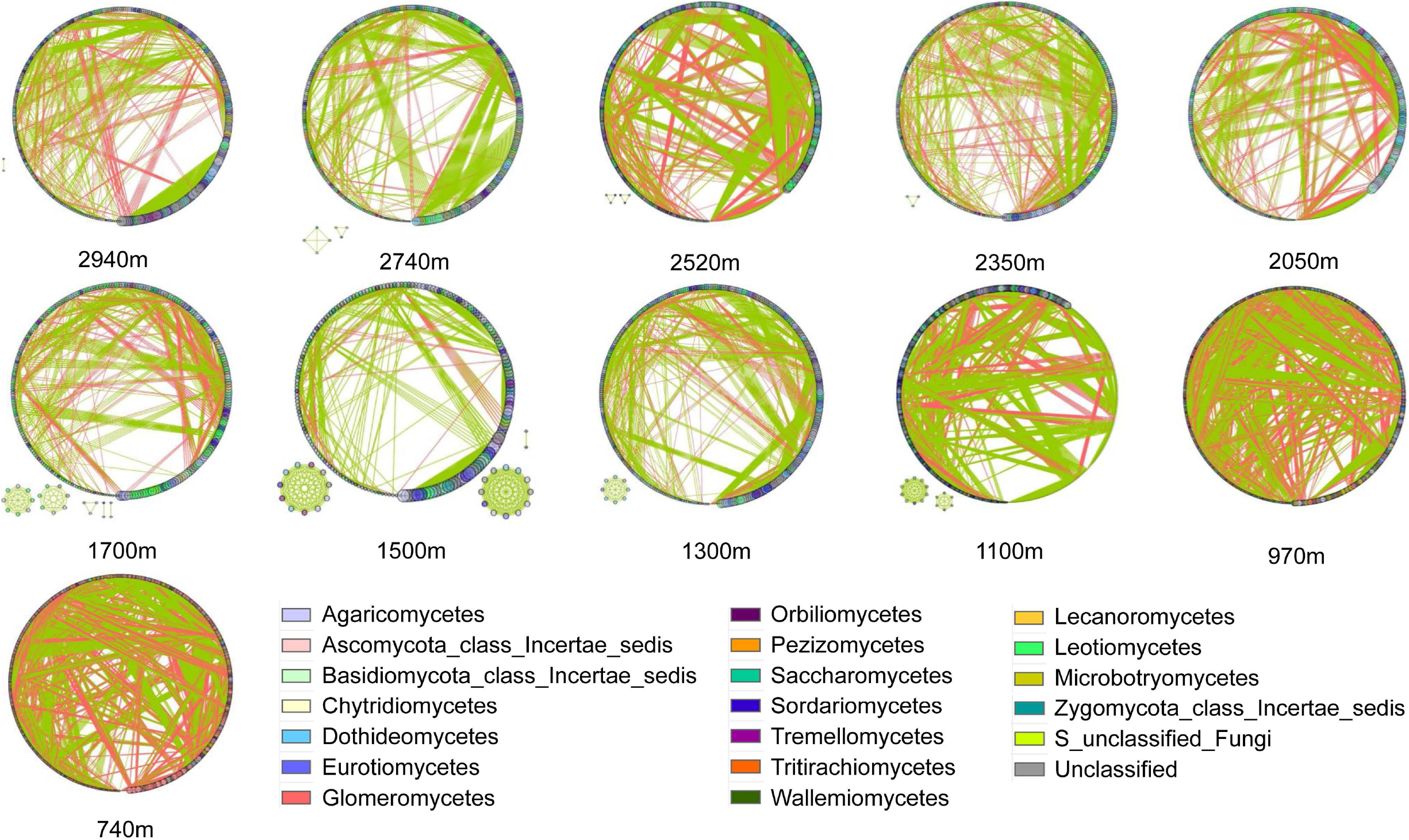
The co-occurrence network connected graph of each elevation gradient. The connection stands for a strong (Spearman’s ρ>0.6) and significant (P-value<0.05) correlation. The nodes each represent unique OTUs in the data sets. The size of each node is proportional to the number of connections (that is, degree). OTUs are coloured by different classes. The green side is positive correlation and the red side is negative correlation.

In comparing the sub-networks of different elevations, for the whole fungal community and each phylum, the network structure of sites at the lower elevations (740–1500 m) was significantly (P <0.05) different from at higher elevations (1700–2940m) (Table S2, S3, S4 and S5 at [https://doi.org/10.6084/m9.figshare.13625741.v1]). Each category of network topological properties of total fungal community exhibited significant elevational patterns (Fig 2). Edge density and betweenness centralization showed unimodal trends, with the count peaking at mid-elevations and lower at both ends of the elevational gradient. Clustering coefficient increased with increasing elevation. The connectivity-elevation relationship was significant for all fungi groups. Between 700 m and 1500 m, the network connectivity decreases with elevation, while at elevations higher than 1500m, the connectivity does not clearly change with elevation. While there is a slight upturn in the curve at the highest elevations, this may be viewed as an artefact of curve-drawing function as there is no clear upward trend in the data points.

**Fig 2.**
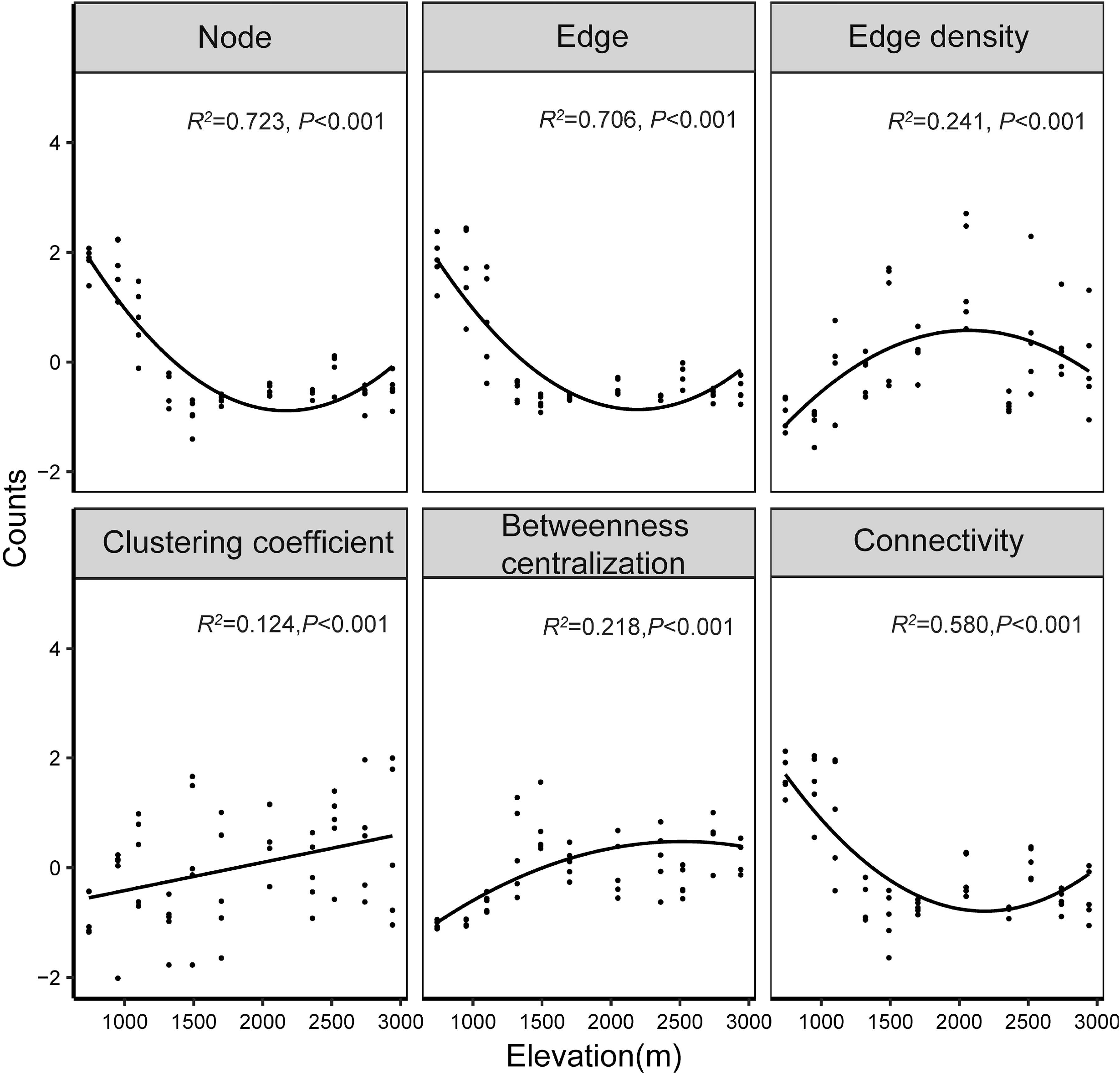
The relationships between network topological properties and elevation for all fungi community. Significant (p<0.001) linear or quadratic models are shown with lines. The better model was selected based on the lower value of Akaike’s information criterion. For better visualization, each network topology feature was z-scale transformed.

The above results suggest that network connectivity at the lower elevations was greater than that of higher, indicating greater network robustness at lower elevations. The main sub-network topological characteristics of each elevation level are shown in Table S6. The number of positive and negative associations decreased with elevation (Figure S4 at [https://doi.org/10.6084/m9.figshare.13625651.v1]), but the proportion of positive correlations among nodes reaches its maximum at 1500 m elevation, and is less above and below this elevation, and the negative correlations are the opposite. For all three phyla that dominated the fungal community - Ascomycota, Basidiomycota and Zygomycota - trends in sub-network connectivity with elevation were roughly the same: each showed a trend of decreasing connectivity with elevation between about 700 m and 1500 m, followed by relatively constant connectivity between 1500 m and 3000 m (Figure S5), this indicated that the network became more discrete and sparser with increasing elevation.

### 2.3 Environment factors influencing the microbial co-occurrence patterns

Based on Random Forest Analysis, MAT and MAP were the major determinants of network connectivity (Fig 3a). Among soil factors, pH, NO_3_^−^-N and NH_4_^+^-N were important drivers of network connectivity (Fig 3b). For each of the three phyla with the highest abundance in the fungal community (Ascomycota, Basidiomycota and Zygomycota), climatic factors (include MAP and MAT) had a strong influence on network connectivity, but other factors such as TC, NO_3_^−^-N, pH and NH_4_^+^-N variously influenced the individual phyla (Figure S6 at [https://doi.org/10.6084/m9.figshare.13625672.v1]). Moreover, microbial diversity has been widely used to determine the influence of biotic factors on the microbial co-occurrence patterns (31), and we found that increased network connectivity was significant correlated with greater alpha diversity (Figure S7 at [https://doi.org/10.6084/m9.figshare.13625681.v1]).

**Fig 3.**
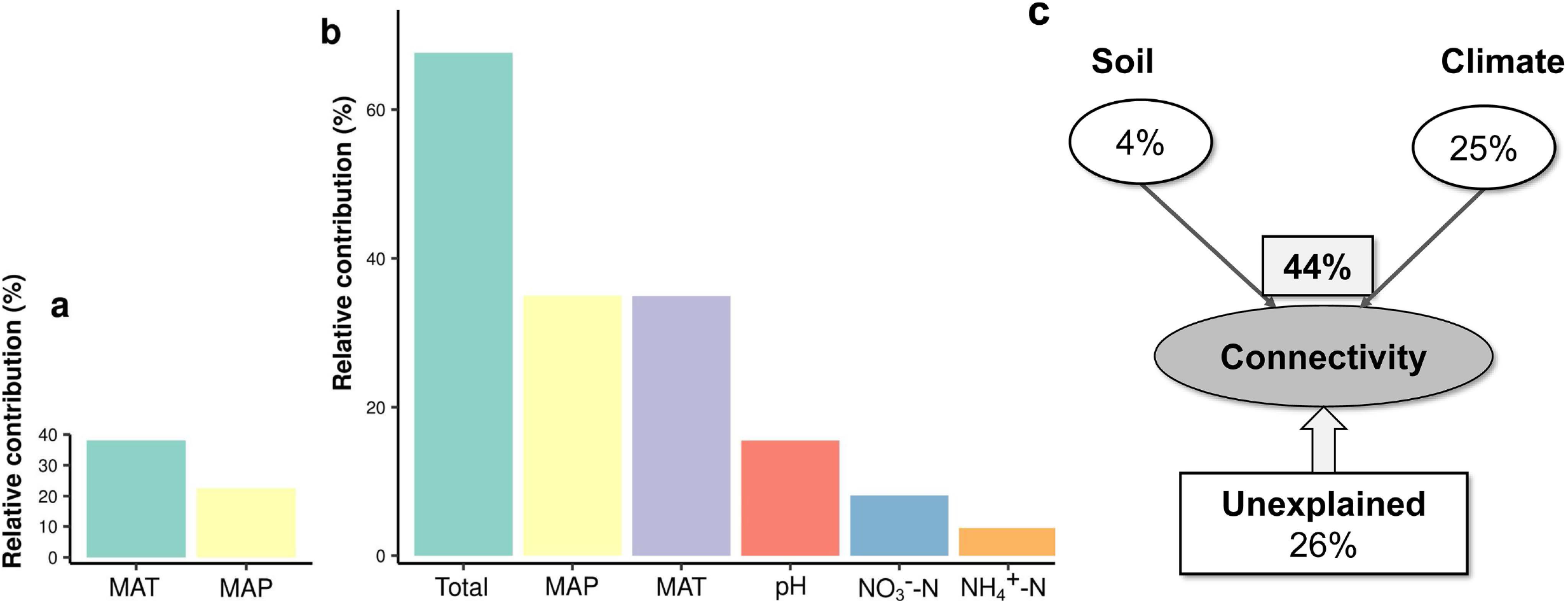
The relative contribution of environmental variables to network connectivity. Pure effects of climate (a), all environmental variables (b) based on random forest analysis, and the contribution of climate and soil parameters calculated based on Variance Partitioning Analysis (VPA) (c). Variation was partitioned by climate factors (MAT and MAP) and soil factors (pH, TC, TN, K^+^, P2O5, Soil texture, NH_4_^+^-N and NO_3_^−^-N). Adjusted R^2^ values are shown here.

VPA was used to estimate the importance of environmental factors for the variation of network structure (32). The pure effects of climate accounted for most of the explained variation of total fungal community network connectivity (25%), and the joint effects of climate and soil variables (44%) captured the main fraction of the explained variation of network connectivity (Fig 3c). The proportion of unexplained variation (adjusted R^2^) was 26%, with the amount explained varying between the three major phyla (Fig 4). Climate together with soil parameters explained varying proportions of the total variation in network properties, with climate factors less important than soil parameters for Zygomycota.

**Fig 4.**
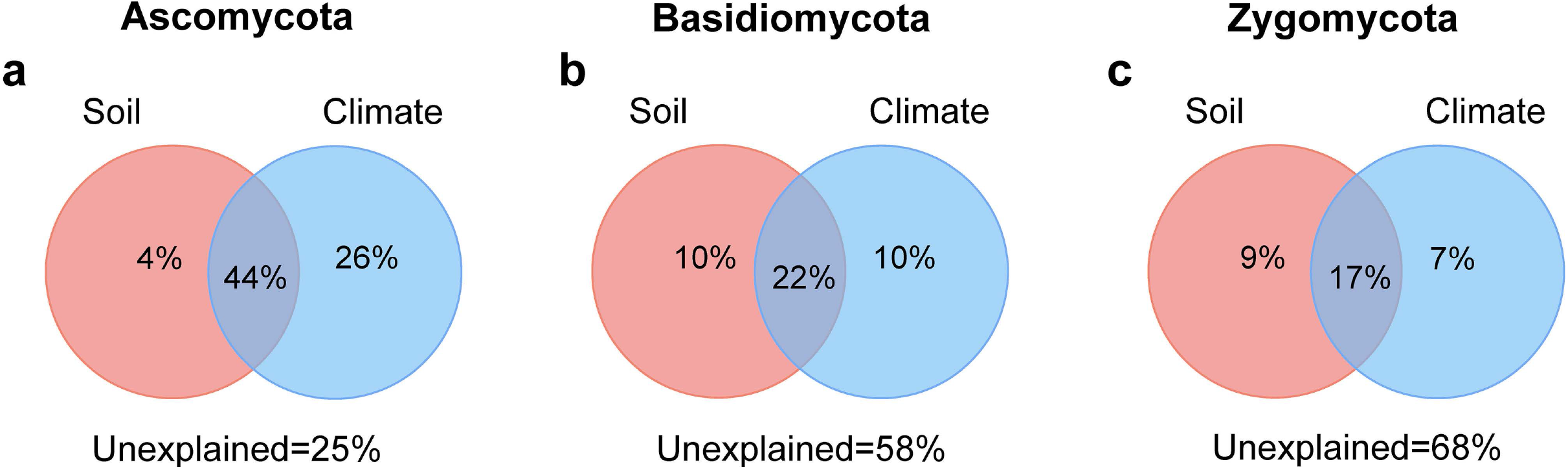
Variation partitioning of network structure by soil physicochemical properties and climate variables (a, Ascomycota; b, Basidiomycota; c, Zygomycota). The specific factors included in each classification were consistent with those in Fig 3. The statistical significance was determined according to Monte Carlo permutation test (9,999, P < 0.01). The values are the adjusted coefficient of determination (adjusted R^2^).

The SEM analysis indicated that the fungal community network connectivity was shaped by hierarchically structured factors, connected to each other by causal relationships (Figure S8). For total fungi community, MAP and MAT apparently affected connectivity directly, and also exerted indirect effects via soil variables, especially pH, and TC. As can be seen from the path coefficient, MAP and MAT were the main influencing factors. The final model explained 71.5% of the variation in network connectivity. For Ascomycota (Figure S9a at [https://doi.org/10.6084/m9.figshare.13625690.v1]), Basidiomycota (Figure S9b) and Zygomycota (Figure S9c) the outcome was broadly similar to that of the whole fungi community, with a dominant role of climate, acting through soil factors. The results are consistent with the analysis of variance partitioning (VPA).

### 2.4 Elevation patterns and influencing factors of fungal trophic guilds

As an additional context to whatever trends occurred in community connectivity, we analysed the contribution of different trophic guilds using FUNguild. Seven recognized fungal trophic types were found on Norikura: symbiotroph, saprotroph-symbiotroph, saprotroph-pathotroph, saprotroph, pathotroph-symbiotroph, pathotroph, and pathotroph-saprotroph-symbiotroph. The ‘unclassified’ trophic mode category was eliminated from the analysis. The relative abundance of trophic guilds differed along the elevational gradient (Figure S10 at [https://doi.org/10.6084/m9.figshare.13625696.v1]). In both the low and high elevations, pathotroph-saprotroph, pathotroph and saprotroph were the the majority while in the mid-elevations the symbiotrophic mode was most abundant.

The three fungal trophic categories with the highest abundance were selected: symbiotroph, saprotroph, pathotroph. The relative abundance of pathotrophs was mainly affected by MAT, and also correlated with TN, NH_4_^+^-N and TC. The relative abundances of saprotrophs were significantly correlated with soil pH, soil texture and TC. For symbiotroph, abundances were controlled by MAP and pH (Figure S11 at [https://doi.org/10.6084/m9.figshare.13625699.v1]). Then, we explored the factors that influence the network connectivity of the three trophic categories. The SEM model showed that the network connectivity of pathotroph was only affected by MAT and MAP. For saprotroph, connectivity is directly affected by MAP, pH and NO_3_^−^-N, and the network connectivity of symbiotic fungi is mainly dominated by NH_4_^+^-N and MAT. The SEM model for saprotrophs explained the greatest proportion of variation in connectivity, at 77.7% (Fig 5).

**Fig 5.**
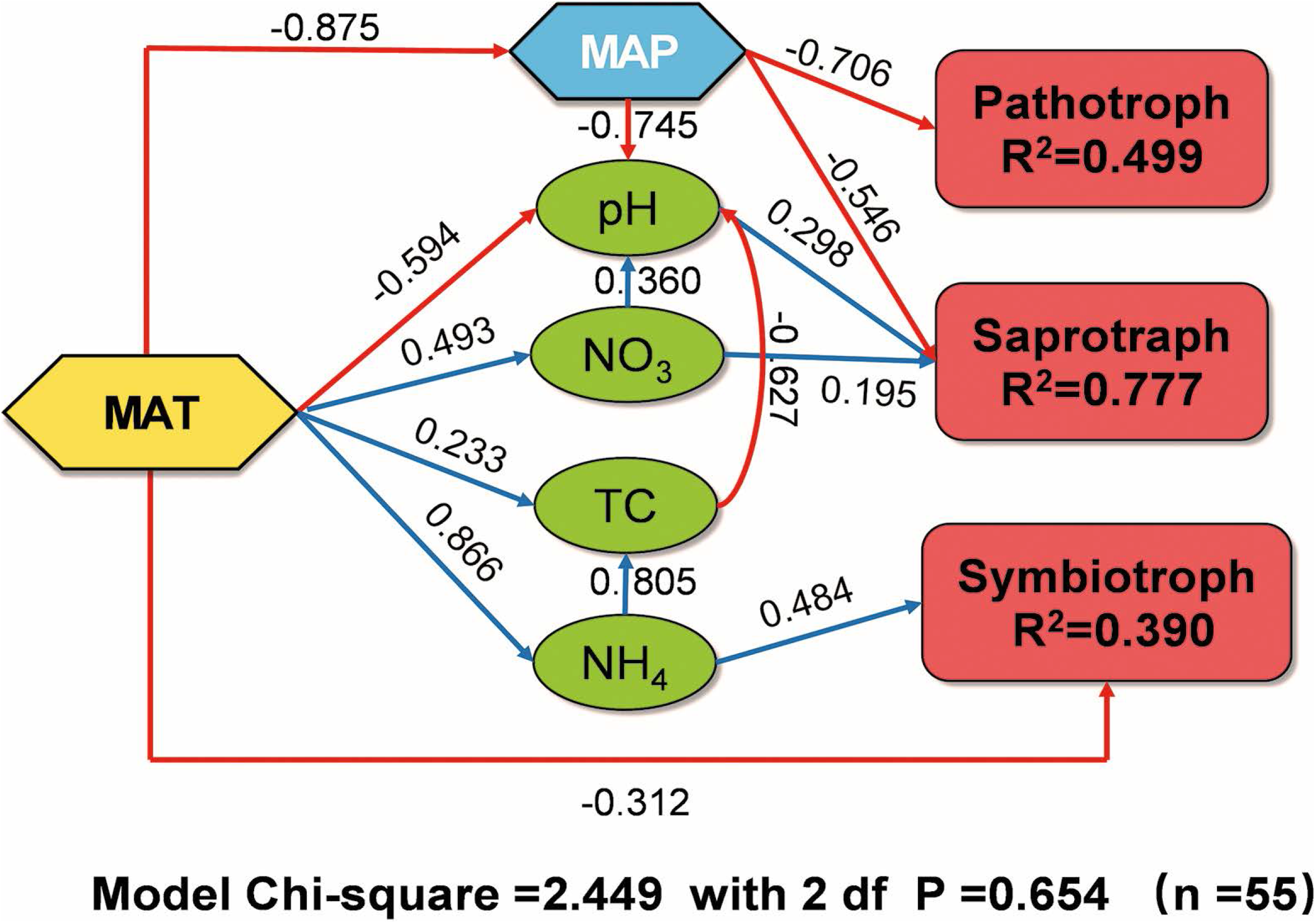
Structural equation model (SEM) in explaining the contribution of environment variables to the network connectivity of symbiotroph, saprotroph and pathotroph soil fungi on Norikura. Blue line: positive correlation; red line: negative correlation. Numbers above/below the arrow lines are indicative of the correlations. The proportion of explained variance (R^2^) appears alongside connectivity in the model. df, degrees of freedom; P, probability level. Significance levels of each predictor are P < 0.05.

### 2.5 Keystone taxa within the fungal community

In the fungal meta-community of Norikura (with all elevations combined) (Fig 6), peripherals accounted for 84.9% of the total nodes, connectors for 13.5% and module hubs for 1.6%, while there were no network hubs, indicating that most of the nodes had only a few links and mostly linked only to the nodes within their own modules. The keystone taxa components are shown in Table S7. In addition, the richness of total keystone taxa was negatively correlated with elevation and MAP (Figure S12 a, b). As for the specific categories, both the abundance of connectors and module hubs showed a decreasing trend with increasing elevation. However, the slope for connectors was −0.005/m, and that for module hubs was −0.01/m, indicating that the decline trend of module hub was slightly faster than that of connector (Figure S12 c). The species composition of the module hubs and connectors varied with elevation (Figure S13 at [https://doi.org/10.6084/m9.figshare.13625705.v1]). For the module hubs, the species richness of high-elevation areas was clearly less than in low elevation areas. At high elevations, *Ascomycota_class_Incertae_sedis* and *Leotiomycetes* dominated, while at low elevations, the proportion of *Sordariomycetes* was larger. For connectors, the trend of species number with elevation was basically consistent with the trend of network connectivity, that is, it decreased at 700-1500 m and then basically remained stable. Each class was uniformly distributed at all elevations, and a single OTU of *Wallemiomycetes* is found at the low elevations.

**Fig 6.**
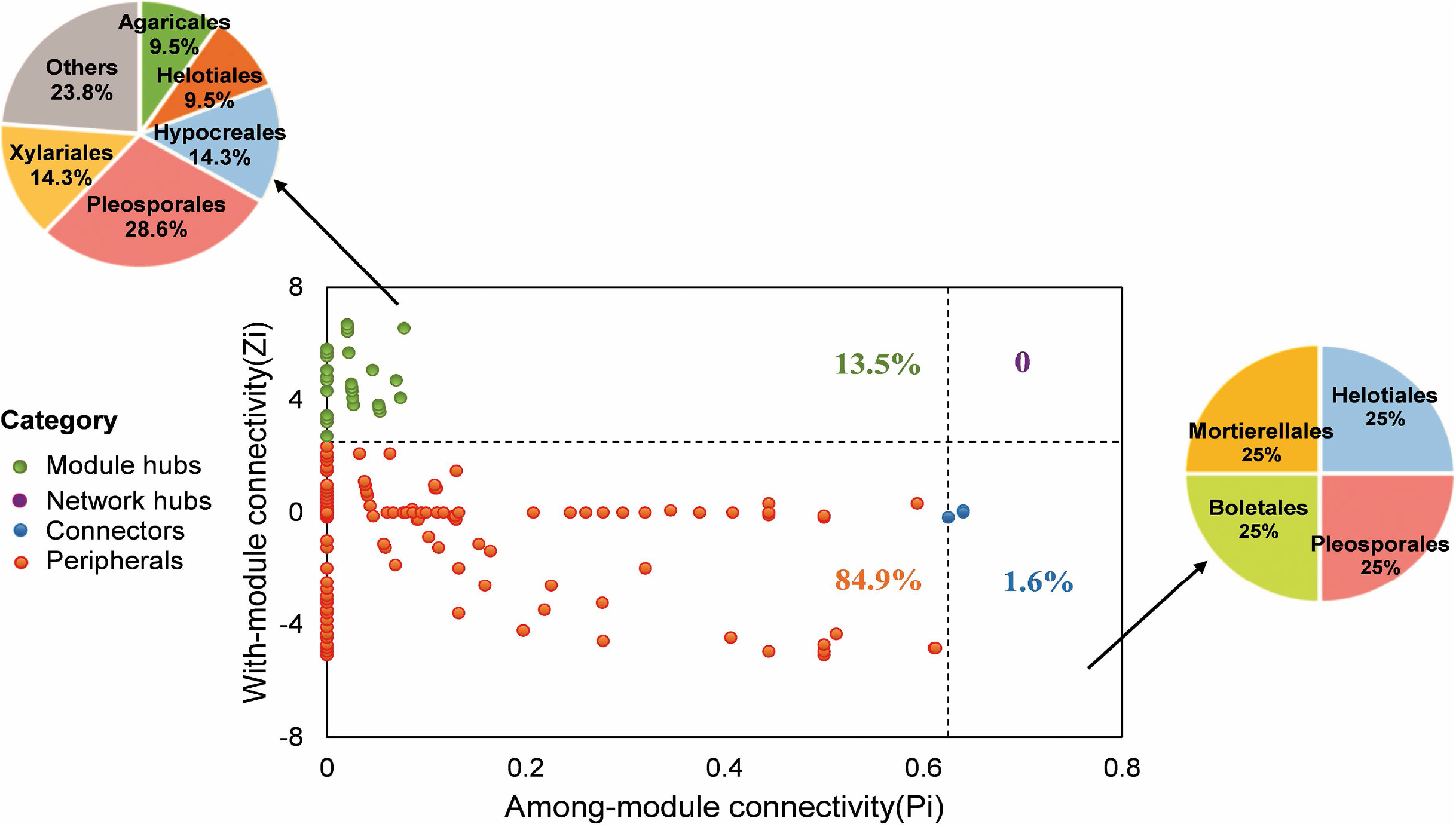
Network roles of analysing module feature at OTU level for total fungi community with the composition of connectors and module hubs. The dots with different colours represent the role of OTU in the network. The pie chart shows the composition of connectors and module hubs respectively, coloured according to Order.

### 2.6 Environmental factors influencing keystone taxa

We constructed additional models by correlating the relative abundance of keystone taxa with environmental properties. Soil pH, MAP and MAT were the variables most closely correlated with the richness of keystone taxa (Figure S14). The SEM model indicated that the factors directly affecting the abundance of keystone taxa included MAP, pH and NH_4_^+^-N. MAT indirectly affected abundance through soil characteristics, and the final model explained 56.9% of the variation. In terms of the association of individual species with environmental variables, higher precipitation inhibited the abundance of keystone taxa, while temperature, pH, and NO_3_^−^-N promoted it. Furthermore, module hubs were more sensitive to environmental changes than the connectors (Figure S15 at [https://doi.org/10.6084/m9.figshare.13625708.v1]).

## 3. Discussion

In our study, our main hypothesis was that as a result of reduced energy flow through the ecosystem, and greater environmental instability, there would be reduced network connectivity within the fungal community at higher elevations, correlating strongly with elevation induced climate change. We also hypothesized a lower diversity of ‘keystone’ taxa at higher elevations due to the prevalence of more generalized interactions.

### 3.1 The complexity of fungal assemblage decreased at low elevation and then stabilized with the rise of elevation

Network analysis showed clear trends in network connectivity of the fungal community with elevation on Mt Norikura. While the overall trend on Norikura agreed with our hypothesis, this was not the steady decrease that would have been expected from the predicted elevation trend in temperature. Instead, for the community of all fungi - and for each of the Basidiomycota, Ascomycota and Zoomycota separately - there was a steep decline in connectivity from 700 m to 1500 m, followed by stabilization between 1500 m and 3000 m.

Fungal communities tend to be tightly associated with plant communities, such as plant community diversity and composition (33, 34, 35). Generally, fungal community connectivity was expected to increase along with plant productivity, based on environmental energy theory (36), according to the principle that more productivity will facilitate the coexistence of more fungal species linked to one another or to other organisms in specialised niches, or narrower niches in which negative associations occur due to competitive exclusion.

Previous studies of Mt Norikura showed that tree diversity decreased with increasing altitude (29), and this might also contribute to the trend of decreasing fungal network connectivity as a result of fewer different types of hosts for mutualism, commensalism, parasitism or decay linking fungal species. On the other hand, the trend might perhaps be a result of greater environmental heterogeneity – for example, soil properties in high elevation areas are generally regarded as less stable, while soil development processes and recovery from disturbance may be slower, while disturbance to vegetation may be more frequent due to avalanches and greater wind speeds. As a result, niche differentitation at lower elevations may be much weaker than those at higher elevations (37). The weaker the niche differentiation, the more generalized the microbial interactions would be, and the fewer network connections expected (5, 22). The observed break point in fungal network structure at 1500m may be determined by the effects of vegetation on soil properties (38, 39, 40, 41), for example caused by the switch to montane boreal-type conifer forest which occurs at this elevation (29), and a shift to abundant ferns (such as *Sasa senanensis*) above 1500m (Fig S16).

### 3.2 Climate as a driver of network variation

The SEM and Random Forest analyses (Fig 3, 4 and S8) support the view that both climate and soil factors (e.g., MAT, MAP and TC) shaped fungal network characteristics, while climate factors played more important roles than soil factors in meta-fungal network assembly.

Both temperature and precipitation had significant effects on network properties on Mt Norikura. Temperature is known to have a major effect on fungal community processes: Decreasing temperature with increasing elevation tends to inhibit soil processes - such as decomposition, nutrient cycling and carbon sequestration - that reducing the diversity and interaction of fungal microorganisms (42). Numerous studies have found that precipitation pattern can play a key role in shaping microbial community structure (43, 44, 45). Wang et al (46) have found that microbial networks became more complex with increasing precipitation in drought-stressed environments, suggesting that this was due to greater nutrient diffusion in water-limited areas, resulting in high plant richness and biomass (47), in turn affecting nutrient supply to soil microorganisms (48). However, our results suggested that greater precipitation resulted in decreased network connectivity, possibly due to excessive soil moisture affecting fungal ecology through waterlogging and restricted oxygen content, leaching or mobility of ions, and the activity of soil animals which turn over soil and physically break up and move litter. Soil waterlogging and lack of oxygen may reduce the potential energy budget for niche specialisation, and more generalized niches should tend not to show up as strong network linkages (5). This is in agreement with the increased negative edge proportion at the higher altitudes (Table 2), where the precipitation stress is more pronounced and such negative relationships could be enhanced.

As for sub-networks, the various microbial phyla responded differentially to soil-related and climate-related factors among habitats (Fig 3 and S9), but in all cases the results indicated that NH_4_^+^-N was a major driver directly affecting the fungal community on the elevation gradients. This was in accordance with the widely accepted view that nitrogen is a particularly important abiotic soil factor affecting soil microbial community (49, 50).

Additionally, we found that the drivers of network complexity were different for different fungal trophic guilds (Fig 5). For pathotrophs, higher precipitation decreased the probability and degree of interaction within the community. Network connectivity of saprotroph fungi community was mainly controlled by soil pH and NO_3_^−^-N, which could be expected as they would affect the supply of substrates by affecting microbial enzymatic activities (51) and changing carbon and nutrient pools in soil environments (52). Temperature and NH_4_^+^-N had the greatest effect on symbiotic fungal communities, possibly by altering plant primary productivity and soil fertility.

These results suggested that the combined role of MAP and MAT did not appear to be equally as important across the different fungal phyla and guilds, apparently reflecting differences in detailed ecology of these groups. However, similar trends obtained from SEM analysis indicate that MAP and MAT play a less direct role but are nevertheless the fundamental drivers.

### 3.3 Structural changes and influencing factors of keystone taxa

Connectivity within and among modules were used to identify the roles of each node in the network (53). Usually, one of four ecological roles (peripherals, module hubs, network hubs or connectors) could be assigned to each node. Topologically, network hubs and connectors represent the regulators or adaptors. Module hubs can be regarded as key elements within distinct modules, which may perform important functions but tend to function at a lower level within the overall community (54).

These keystone taxa have also vital ecological functions in the microbial community. In the present study, we did not detect network hubs. The fungal module hubs belonging to *Agaricomycetes* (55), *Dothideomycetes* (56), and *Leotiomycetes* (57) have the potential to improve nutrient acquisition and combat pathogenic taxa, and maintain cooperative metabolic associations with other species. While the connector species belonging to *Sordariomycetes* play an important role in ecosystems and some of them have the potential to produce bioactive compounds (58), In general, network hubs, module hubs and connectors had diverse metabolisms.

In our study, more keystone taxa occurred at lower altitudes (Fig S12), which might be explained, in part, by the increasing biomass stimulated at low elevations where plant diversity is high, providing more opportunities for different species to interact with each other (59). This also supports the conclusion that low altitude region has more complex network structure. In addition, some of the keystone taxa are closely related to environmental factors, such as *Sordariomycetes,* the class includes many important plant pathogens, as well as endophytes, saprobes, epiphytes, coprophilous and fungicolous, lichenized or lichenicolous taxa (60), which are more sensitive to environmental changes (Fig S15).

### 3.4 Is elevation an analogue of latitude?

The finding that MAP variation with elevation has a key role in variation in fungal network structure on a mountain elevation gradient was surprising, as in mesic climates temperature is usually seen as the key variable affecting elevation gradients (61). There is a widespread paradigm that elevation gradients may be seen as an analogue of latitudinal temperature gradients, but in miniature (61). However, as the example of this study shows, the analogy may be too simplistic. Latitudinal gradients in temperature in mesic regions do not typically show a peak in precipitation in boreal latitudes but instead a steady decline in precipitation towards high latitudes (e.g. the eastern sides of North America and Asia).

## 4. Conclusions

This study provides evidence of a decrease in soil fungal network connectivity towards higher elevations of mid latitude mountains, with both mean annual precipitation (MAP) and mean annual temperature (MAT) playing important underlying roles, and the effects of climate on soil factors being important in this. This trend is presumably associated with broader niches and less specific associations at higher elevations. Limited data on the trend in tree species diversity in Norikura suggests that this might also play a role.

The importance of variation in precipitation rather than temperature alone suggests that latitudinal trends in connectivity may not resemble elevation trends, and should be considered separately. It would be intriguing to compare the trends observed here with other long mountain elevation series, and long latitudinal series which also cross a wide range of biome types, to discern the general rules which structure network connectivity in fungal communities.

## 5. Materials and Methods

### 5.1 Site description and sampling

Mt. Norikura is an extinct volcano (last active about 18,000 years ago) located at the border of Gifu and Nagano prefectures in Central Japan, reaching ~3026 m above sea level (62). It has a cool temperate monsoon climate in a climate station at its base at 1000 m a.s.l., with a mean annual temperature (MAT) of 8.5 °C, and a mean annual precipitation of ~2206 mm. Extrapolating according to the moist air lapse rate, the total mean annual temperature gradient is expected to be between 11 ℃ and −4 ℃ (29).

The whole of the mountain from slightly below 700 m upwards has a cover of late Quaternary volcanics, while below 700 m it is underlain by Quaternary granite (29). The soil supports a mixed vegetation comprised of montane deciduous broad-leaved forest zone at low elevation (800-1600 m), a subalpine coniferous forest zone in mid-elevations (1600-2500 m) and open pine scrub at high elevations (2500-3000 m; 29). This silicon-rich andesite mixture and humus make the soil acidic, and the pH increases slightly with elevation. The vegetation cover above 1500 m is almost undisturbed by humans, except in some areas right at the summit (we avoided sampling these areas). As the whole area is protected as a national park, it is in a relatively pristine state.

Sampling was carried out along a transect on the eastern slope of the mountain from late July to early August (in 2015), collecting a total of 55 soil samples from 11 elevational isoclines, each separated by ~200 m of elevation. To eliminate the effect of high spatial heterogeneity on microbial analyses, five separate composite soil samples were taken at each elevational level (spaced 100 m apart), each sample consisting of a composite of five cores taken within a 10 m x 10 m square: one at each corner and one in the centre. Each core was approximately 5 cm in diameter and 10 cm deep, consisting only of the about 10 cm of soil in the lower part of A horizon (defined as having at least some mineral grains present) – any leaf litter, O horizon and the upper part of A (pure organic) horizon was removed before sampling (63). The soil sample collection diagram was shown in Fig S17 at [https://doi.org/10.6084/m9.figshare.13625714.v1]. Soil samples were transported to the laboratory in an ice cooler to minimize postharvest changes in biota. Soil samples were sieved using 2 mm mesh to remove roots and stones, homogenized, and stored at 4 °C for soil physicochemical measurements and at −20 °C for DNA extraction.

### 5.2 Soil chemical properties and climate data

Soil pH, texture, total carbon (TC), total nitrogen (TN), P_2_O_5_, NO_3_^−^-N and NH_4_^+^-N and K^+^ were measured in each sample at Shinshu University, using standard Soil Science Society of America (SSSA) protocols. Soil pH was measured in a soil distilled water suspension (Kalra 1995). Soil were stored after drying soil samples at room temperature. Total carbon and nitrogen contents were measured by using an elemental analyser (Flash EA 1112, Thermo Quest Ltd., USA). Concentrations of PO_4_^−^, NH_4_^+^, NO_3_^−^ and K^+^ were determined using a reflectometer (Merck Ltd., Germany). Concentrations of PO4−, NH4+ and NO3− were converted to equivalent P2O5, NO_3_^−^-N and NH_4_^+^-N, respectively.)

Climate data were derived using the orographic model of Land, Infrastructure, Transport and Tourism (http://nlftp.mlit.go.jb/ksj/gml/datalist/KsjTmplt-G02.html) to derive mean annual temperature and precipitation. This model utilizes input on topography, lapse rate, geographical position relative to the coastlines, and wind direction, interpolated from weather stations, to give mean annual temperature (MAT) and mean annual precipitation (MAP) surfaces for Japan (64).

### 5.3 High throughput sequencing

DNA was extracted from 0.5 g of soil using a Power Soil DNA extraction kit (MoBio Laboratories, Carlsbad, CA, USA) following protocol described by the manufacturer. Concentration and quality of extracted DNA was determined with spectrometry absorbance between 230–280 nm detected by a NanoDrop ND-1000 Spectrophotometer (NanoDrop Technologies) and OPTIMA fluorescence plate reader (BMG LABTECH, Jena, Germany). Fungal DNA were subsequently amplified by PCR targeting the internal transcribed spacer (ITS2) region with the primer combination, ITS86F (5′-GTGAATCATCGAATCTTTGAA-3′) and ITS4(R) = (5′-TCC TCCGCTTATTGATATGC-3′). PCR was performed in 50 μl reactions using the following conditions: 95 °C for 10 mins; 30 cycles of 95 °C for 30 s, 55 ℃ for 30 s, 72 °C for 30 s and 72 °C for 7 min. PCR products were purified using the QIAquick PCR purification kit (Qiagen) and quantified using PicoGreen (Invitrogen) spectrofluorometrically (TBS 380, Turner Biosystems, Inc. Sunnyvale, CA, USA). ITS Sequencing was done using Illumina Miseq platform (Illumina, Inc., San Diego, CA, USA) at the Center for Comparative Genomics and Evolutionary Bioinformatics, Dalhousie University, Canada according to protocols enumerated in Op De Beeck et al., Comeau et al. (64)

### 5.4 Sequence processing

The raw ITS reads were obtained from the Miseq sequencing machine in fastq format. Sequence data was then processed using Mothur (version 1.32.1, http://www.mothur.org) following the Mothur Miseq SOP. Forward and reverse directions, which were generated as separated files were combined using the make.contiq command. Sequences with lengths less than 200 bp were removed using the screen. seqs command (65, 66). Putative chimeric sequences were detected and removed via the Chimera Uchime algorithm contained within Mothur in de novo mode. Rare sequences (less than 10 reads) were removed to avoid the risk of including spurious reads generated by sequencing errors. High quality sequences were assigned to OTUs (operational taxonomic units) at ≥99% similarity. Taxonomic classification of each OTU was done using classify command in mother at 80% naïve Bayesian bootstrap cut-off with 1000 iterations against the UNITE database. The final OTU table consisted of 476590 sequences (average of 8665 sequences per sample) distributed into 16382 OTUs, of those 7284 were represented by more than 1 sequence. All the sequences used and their information have been deposited in the National Center for Biotechnology Information Sequence Read Archive (accession code SRP140430). (64)

### 5.5 Statistical analysis

The co-occurrence network was constructed with the ‘WGCNA’ R package based on the Spearman correlation matrix, which provides a comprehensive and flexible set of functions for performing weighted correlation network analysis. (67). For whole fungi community, we kept OTUs with relative abundances greater than 0.01% for fungal communities (22), only OTUs occurring in more than 40% of all samples were kept for network construction (68). Based on correlation coefficients and the Benjamini and Hochberg false discovery rate (FDR) adjusted P-values for correlation, which was implemented in the ‘multtest’R package (69). The cutoff of FDR-adjusted P-values was 0.001. The nodes and the edges in the network represent OTUs and the correlations between pairs of OTUs, respectively. To reduce network complexity and facilitate the identification of the core soil community, correlation coefficients (r) with an absolute value > 0.60 and statistically significant (P < 0.05) were used in network analyses (Barberán et al., 2012). Gephi (https://github.com/gephi) was used to generate the network image (70).

A set of network topological properties (number of positive correlations, number of negative correlations, edge density, clustering coefficient, betweenness centralization, and connectivity) was calculated for each co-occurrence network with the ‘igraph’ package in R (71). Among these indexes, connectivity represents the number of edges connected to a node, clustering coefficient reflects the higher connectedness among nodes in a particular region of a network (72), betweenness centrality reveals the role of a node as a bridge between components of a network. Permutation multivariate analysis of variance (PERMANOVA, also known as Adonis) based on Euclidean distance was further applied to examine the differences in the topology structure of various elevation gradients, using the function “adonis” of the vegan package v 2.4.6 in R v3.4.3, and the entire (non-subsampled) dataset (73). We used the Wilcoxon rank-sum test to determine the difference in the network-level topological features between different elevations. In order to assess the network topological properties elevation pattern of fungal community, based on Akaike’s information criterion (74) and the function stepAIC with backward stepwise model selection in the R package MASS, we performed the most suitable linear or quadratic regression fitting model.

Keystone taxa have been identified by measuring the degree of association with other species within co-occurrence networks (14). We calculated within-module connectivity (Zi) and among-module connectivity (Pi) of each node based on Markov cluster algorithm with the ‘rJava’ package in R. The threshold values of Zi (which describes how well a node is connected to other nodes within its own module) and Pi (which reflects how well a node connects to different modules) for categorizing OTUs were 2.5 and 0.62, respectively (75). Here we define nodes as network hubs (Zi > 2.5; Pi > 0.62), module hubs (Zi > 2.5; Pi < 0.62), connectors (Zi < 2.5; Pi > 0.62) and peripherals (Zi < 2.5; Pi < 0.62), based on their within-module degree (Zi) and participation coefficient (Pi) threshold value (76, 77), which could determine how the node is positioned within a specific module or how it interacts with other modules (78). Network hubs were highly connected, both in general and within a module, the module hubs were highly connected within a module, the connectors provided links among multiple modules, and the peripherals had few links to other species. Network hubs, module hubs, and connectors were termed keystone network topological features; these are considered to play important roles in maintaining community stability and resisting environmental stress (79); thus, we define the OTUs associated with these nodes as keystone taxa (80). We used Spearman’s correlation to check the relationships between keystone taxa richness and the ecological factors.

Ecological guilds based on trophic mode were assigned using the FUNguild (http://www.stbates.org/guilds/app.php). This tool was able to assign trophic mode and guild to fungal taxa, based on comparison to a curated database of fungal life styles and use of resources. Trophic mode refers to the mechanisms through which organisms obtain resources, hence potentially providing information on the ecology of such organisms (81). The relative abundance of the FUNguild results (trophic mode) were used to interpret the communal and taxa roles at each elevation. Only sequence taxonomy identity above 93% and the guild confidence ranking assigned to ‘highly probable’ and ‘probable’ was accepted.

Since MAT was calculated by using mean lapse rate of 0.6 °C/100 m, it showed a complete correlation with elevation and is used to represent the effect of elevation. We used the remaining environmental variables for further analysis (MAP, pH, TC, TN, K^+^, P2O5, soil texture, NH_4_^+^-N and NO_3_^−^-N). To check relationship between network topological features and environmental properties, raw environmental data were standardized to make the different environmental factors comparable. We performed random forest analysis to explore the contribution of environmental factors to network connectivity.

Random Forest is a powerful machine learning tool that offers high prediction accuracy by using an ensemble of decision trees based on bootstrapped samples from a dataset (82). It was performed with 999 permutations using the ‘randomForest’ and ‘rfPermute’ packages. The best predictors were identified based on their importance using the importance and varImpPlot functions. Increase in node purity and mean square error values were used to determine the significance of the predictors using the random Forest Explainer package (83). The factors significant at P< 0.01 were selected as the predictors of network connectivity. To ensure that overall results were independent of the chosen method, we carried out variation partitioning (84) to partition the variation in the response variable with respect to the soil variables (pH, TC, TN, K^+^, P2O5, Soil texture, NH_4_^+^-N and NO_3_^−^-N), climate (MAT, MAP) and their joint effects. Variation partitioning was conducted using the varpart function in the R package vegan (85).

Finally, to examine the causal relationships between network topological features and elevation/soil chemical properties, we constructed structural equation models (SEM). We first constructed an initial model for each taxonomic group that included all possible pathways between the response variable, the key soil chemical variables and elevation. In addition to direct pathways, we considered indirect ones to see if variables that were not directly related to the response variable exerted some effects via other mediating variables. All variables were standardized before they were entered in the SEM. From the initial model, non-significant paths were eliminated stepwise until all remaining paths were significant (if possible) and directly or indirectly related to the response variable. The goodness of fit of the final model was evaluated with a chi-square test; a non-significant p-value (>0.05) indicates that there are no significant deviations between the model and data (86).

## Acknowledgements

This work was partly supported by a National Research Foundation (NRF) grant funded by the Korean Government, Ministry of Education, Science and Technology (MEST) (NRF-0409– 20150076). And YS’s work was supported by the National Natural Science Foundation of China (42077053). JA acknowledges the generous assistance of Nanjing University and the School of Geography and Oceanography in supporting this research.

## Tables

Table S1. The number of associations between fungal OTUs at the order level.

Table S6. Key topological features of each elevation gradient.

Table S7. Keystone taxa composition of the whole fungal community.

---

Figure S1. Environmental variables shown in relation to elevation.

MAP: Mean annual precipitation; MAT: Mean annual temperature; TC: total carbon; TN: total nitrogen; NH4^+^-N: ammonium nitrogen; NO_3_^−^-N: nitrate nitrogen; K:potassium. P2O5: available phosphorus. The values of TC and TN are shown in percentages. Total of percentage silt and clay content are used to indicate soil texture. Shown are adjusted R^2^ values.

Figure S2. The co-occurrence network interaction of meta-fungi community of Mt Norikura, at all elevations. Nodes are coloured by different classes. Green is positive correlation; red is negative correlation.

Figure S5. Key network parameters of the three phyla vary with the elevation. The solid line means p- value is less than 0.05, and the dashed line means insignificant.

Figure S8. Structural equation model explaining the contribution of environmental variables to fungal community network connectivity. The values corresponding to the pathways are standardized path coefficients. Blue arrows indicate positive effects, red arrows denote negative effects. R^2^values indicate the amount of explained variations in the response variables.

Figure S12. Plots showing the relationship between total keystone taxa and elevation (a) and precipitation (b), elevation trend of abundance of coonector and module hub (c). Significance level is * P ≤ 0.05, ** P ≤ 0.01 and *** P ≤ 0.001.

Figure S14. The contribution of environmental factors that correlate with the richness of keystone taxa

Figure S16. Altitudinal changes in number of species of trees on Mount Norikura. Soild and open circles represent trees shorter and taller than 1.3m, respectively.

